# Comparison of the full distribution of fitness effects of new amino acid mutations across great apes

**DOI:** 10.1101/696971

**Authors:** David Castellano, Moisès Coll Macià, Paula Tataru, Thomas Bataillon, Kasper Munch

## Abstract

The distribution of fitness effects (DFE) is central to many questions in evolutionary biology. However, little is known about the differences in DFEs between closely related species. We use more than 9,000 coding genes orthologous one-to-one across great apes, gibbons, and macaques to assess the stability of the DFE across great apes. We use the unfolded site frequency spectrum of polymorphic mutations (n = 8 haploid chromosomes per population) to estimate the DFE. We find that the shape of the deleterious DFE is strikingly similar across great apes. We confirm that effective population size (*N_e_*) is a strong predictor of the strength of negative selection, consistent with the Nearly Neutral Theory. However, we also find that the strength of negative selection varies more than expected given the differences in *N_e_* between species. Across species, mean fitness effects of new deleterious mutations co-varies with *N_e_*, consistent with positive epistasis among deleterious mutations. We find that the strength of negative selection for the smallest populations: bonobos and western chimpanzees, is higher than expected given their *N_e_*. This may result from a more efficient purging of strongly deleterious recessive variants in these populations. Forward simulations confirm that these findings are not artifacts of the way we are inferring *N_e_* and DFE parameters. All findings are replicated using only GC-conservative mutations, thereby confirming that GC-biased gene conversion is not affecting our conclusions.

## Introduction

All organisms undergo mutation. Studying the effect of mutations on fitness is fundamental to explain the patterns of genetic diversity within and between species and to predict the impact of population size on the probability of survival, or extinction of a species (Eyre-Walker and Keightley 2007). The fitness effects of all new mutations that can occur in a given genome are described by the distribution of fitness effects (DFE). The allele frequency distributions or the site frequency spectrum (SFS) contains information to infer the DFE of new mutations. Current statistical methods infer the DFE while accounting for demography and other sources of distortion in the SFS (Eyre-Walker *et al*. 2006; Keightley and Eyre-Walker 2007; Boyko *et al*. 2008; Schneider *et al*. 2011; Galtier 2016; Kim *et al*. 2017; Tataru *et al*. 2017; Barton and Zeng 2018).

The DFE of new deleterious amino acid mutations is assumed to be similar in closely related species, while the mean effect of those deleterious mutations (*S_d_* = 2*N_e_s_d_*) is expected to increase as a function of the effective population size (*N_e_*) (Kimura 1983; Ohta 1992). Very little is known about the variability in the DFE of new beneficial mutations across species as well as how this variability affects the estimates of the deleterious DFE. Most studies have described the DFE of new deleterious mutations of individual populations, while the study of the differences in the DFE between populations or species has remained unexplored. To our knowledge, Huber *et al*. (2017) is the first work that tries to explain, under different theoretical models, the detected differences in the deleterious DFE of new amino acid mutations in humans, Drosophila, yeast, and mice. They find that humans have more strongly deleterious mutations than Drosophila and that species complexity (measured by the number of different phenotypic dimensions under stabilizing selection) correlates positively with the degree of deleteriousness of new mutations as expected under Fisher’s Geometrical Model (Fisher 1930). They find that these differences in the DFE cannot be explained by differences in population size or demography between the species. Recently, Zhen *et al*. (2018) argued that there is a higher proportion of beneficial amino acid changing mutations in humans compared to mice and flies when accounting for the population size of the outgroup species. Zhen *et al*. (2018) use divergence to detect beneficial mutations that reached fixation many generations ago, many of those were strongly beneficial mutations that contribute disproportionately more to divergence than polymorphism. Here we focus instead on new weakly beneficial (still segregating) mutations. The power of our approach to infer with precision the tail of the beneficial DFE is reduced. However, relying solely on polymorphisms means that our estimates are more robust to ancient fluctuations in the effective population size that impacted the probability of fixation of slightly selected mutations in the ancestral populations (Rousselle *et al*. 2018).

Whether the full DFE varies on shorter timescales remains an open question. It also remains to be considered the contribution of new weakly beneficial mutations to the SFS and by extension the estimates of the deleterious DFE. Tataru *et al*. (2017) have shown that beneficial mutations can indeed contribute to the SFS and affect the estimates of the deleterious DFE. In fact, Huber *et al*. (2017) reported more weakly beneficial mutations in humans than in *Drosophila* indicating that weakly beneficial mutations are relevant in humans and probably are also important in other non-human great apes. A recent study has found that ∼¾ of adaptive substitutions in the human lineage were driven by weakly beneficial amino acid mutations (Uricchio *et al*. 2019). In this work, we compare the DFE in a panel of closely related species - the nine great ape populations studied by Prado-Martinez *et al*. (2013). Great apes share most of their genes and genomic configuration. Gene density, mutation rate and recombination rate (at least at broad scale) are very similar in these species (Stevison *et al*. 2016; Smith *et al*. 2018; Kronenberg *et al*. 2018; Besenbacher *et al*. 2019). Thus, our inference of the DFE is affected only marginally by variation in these factors. Interestingly, there is substantial variation in the effective population size (*N_e_*) and remarkable differences in the population histories of great apes (Mailund *et al*. 2011; Scally *et al*. 2012; Prado-Martinez *et al*. 2013; McManus *et al*. 2015; Bataillon *et al*. 2015; de Manuel *et al*. 2016). This variation in *N_e_* allows us to test predictions of the Nearly Neutral Theory (Ohta 1992). By polarizing mutations and using the unfolded SFS (uSFS), we are able to infer the DFE of new weakly beneficial mutations without relying on divergence and make a cleaner inference of the current deleterious DFE. This allows us to investigate if *N_e_* also has an impact on the beneficial DFE. For instance, it is commonly assumed that the rate and effect size of beneficial and deleterious mutations are shared between closely related species, but these quantities might change if the fitness of the population changes (Silander *et al*. 2007). Recent works have shown that GC-biased gene conversion (gBGC) can bias the inference of the DFE and population history (Pouyet *et al*. 2018; Bolívar *et al*. 2018). Here we also assess the impact of gBGC by replicating our analyses using only GC-conservative mutations (A<->T and C<->G) which are unaffected by gBGC.

Finally, we use the method of Tataru and Bataillon (2019) (polyDFEv2.0) to test for invariance of DFEs across species. Under this new method any fitted parameter can be shared across species or fitted independently for each species. In this work, we compare the fit of several models to investigate which aspects of the full DFE are shared and which aspects differ between great apes.

## Materials and Methods

### Data Sets

SNP calls from the autosomes are retrieved from Prado-Martinez *et al*. (2013) for nine great ape populations: *Homo sapiens* (this sample includes 3 African and 6 Non-African individuals), *Pan paniscus*, *Pan troglodytes ellioti*, *Pan troglodytes schweinfurthi*, *Pan troglodytes troglodytes*, *Pan troglodytes verus*, *Gorilla gorilla gorilla, Pongo abellii* and *Pongo pygmaeus*. Hereafter we will refer to these species as humans, bonobos, Nigeria-Cameroon chimpanzees, eastern chimpanzees, central chimpanzees, western chimpanzees, western lowland gorillas, Sumatran orangutans and Bornean orangutans, respectively. To allow a fair comparison between species (note that only 4 individuals of western chimpanzees and central chimpanzees were sequenced) we downsample all great apes to 8 randomly chosen haploid chromosomes, while positions called in less than 8 chromosomes are discarded. Supplementary Table 1 shows the final list of analyzed genes per species. In Prado-Martinez *et al*. (2013) all reads are mapped to the human reference genome (hg18). We lift over the original VCF to hg19/GRCh37.75 coordinates to take advantage of more recent functional annotations. To avoid errors introduced by miss-mapping due to paralogous variants and repetitive sequences, we also restrict all analyses to a set of sites with a unique mapping to the human genome as in Cagan *et al*. (2016). Additionally, we require positions to have at least 5-fold coverage in all individuals per species. Only the remaining set of sites are used in further analyses. Genomes are annotated using the SnpEff and SnpSift software (Cingolani *et al*. 2012) (version 4.3m, last accessed June 2017) and the human database GRCh37.75. We extract 0-fold non-synonymous and 4-fold synonymous sites from the codon information for the canonical transcript provided by SnpEff (2-fold and 3-fold degenerate sites are discarded). We assume that the degeneracy and gene annotations are constant across species.

### Polarization of Mutations

To estimate the full DFE using DNA diversity data we need first to polarize SNPs to call derived variants (Boyko *et al*. 2008; Schneider *et al*. 2011; Tataru *et al*. 2017). We use two outgroup species (*Nomascus leucogenys* or gibbons, and *Macaca mulatta* or macaques) and a probabilistic method to polarize SNPs using Kimura 2-parameter (K2) model (Keightley and Jackson 2018). We downloaded a multiple species alignment from the UCSC server between all known coding sequences in the human reference genome (GRCh37.75/hg19), gibbon (Nleu3.0/nomLeu3) and macaque (BGI CR_1.0/rheMac3) (from:http://hgdownload.cse.ucsc.edu/goldenPath/hg19/multiz100way/alignments/knownGene.exonNuc.fa.gz). Sites are retained for analysis if there is no missing data in the focal species or either outgroup species. We also removed CpG sites, as CpG hypermutability may result in polarization errors (Keightley and Jackson 2018). CpG sites are defined as sites that are CpG in their context in either the focal species or any of the outgroup (including both REF and ALT alleles). In doing so, we remove 4% of all 0-fold non-synonymous and 5% of all 4-fold synonymous sites that passed our previous quality filters. We compare the uSFS from the K2 model and a more complex model allowing six symmetrical rates between all potential nucleotide changes (R6 model) obtaining indistinguishable uSFS for synonymous and non-synonymous mutations (data not shown). Supplementary Table 1 shows the uSFS for synonymous and non-synonymous mutations in all great apes. All our results can be recapitulated from that table.

### Estimation of the DFE and bootstrapping

We use the polyDFEv2.0 framework (Tataru and Bataillon 2019) to estimate and compare the DFE across species by means of likelihood ratio tests (LRT). The inference is performed on the uSFS data only (divergence counts to an outgroup are not fitted) and uSFS data is fitted using a DFE model comprising both deleterious (gamma distributed) and beneficial (exponentially distributed) mutations. polyDFE assumes that new mutations in a genomic region arise as a Poisson process with an intensity that is proportional to the length of the region and the mutation rate per nucleotide (*μ*). We assume that *μ* remained constant across great apes (Besenbacher *et al*. 2019). Both an ancestral SNP misidentification error (*ε)* and distortion parameters (*r_i_*) can be estimated. See Table 1 and 2 for the list of parameters estimated in each model. To ensure that the likelihood function is reliably maximized we perform 10 runs of maximization with randomly starting values.

**Table 1.** Parameters included in each model used for estimating the DFE. In grey the two models on which of our results are based. *S_d_* is the mean population scaled selection coefficient of new additive deleterious mutations (s ≤ 0); *b* is the shape parameter of the deleterious DFE; *S_b_* is the mean population scaled selection coefficient of new additive beneficial mutations; *p_b_* is the proportion of new beneficial mutations (s > 0); *r_i_* is the series of nuisance parameters used to control for potential distorters of the uSFS; *ε_anc_* is the polarization error. *θ* is the population scaled mutation rate (*4N_e_μ)*.

| Model | $S_d$ | $b$ | $S_b$ | $p_b$ | $r_i$ | $\epsilon_{anc}$ | $\theta$ |
| --- | --- | --- | --- | --- | --- | --- | --- |
| 0 | fitted | fitted | fitted | fitted | 1 | 0 | fitted |
| 1 | fitted | fitted | fitted | fitted | fitted | 0 | fitted |
| 2 | fitted | fitted | fitted | fitted | fitted | fitted | fitted |
| 3 | fitted | fitted | 0 | 0 | fitted | 0 | fitted |
| 4 | fitted | fitted | 0 | 0 | fitted | fitted | fitted |

The estimation of the DFE entails substantial statistical uncertainty. To obtain the sampling variance of parameter estimates and approximate confidence intervals we use a bootstrap approach. Bootstraps are generated by re-sampling the data at the site level by parametric bootstrapping. We assume that all counts in the uSFS are independent variables following a Poisson distribution, with means specified by the observed uSFS data. This is in line with the modeling assumption that the number of mutations in each uSFS entry follows a Poisson process (Tataru *et al*. 2017). We use the R function *bootstrapData*() (from: https://github.com/paula-tataru/polyDFE/blob/master/postprocessing.R) to get the bootstrap replicates.

### Statistical Test for Differences in DFEs Across Species

polyDFE returns maximum likelihood (ML) estimates and therefore, LRTs and the Akaike information criteria (AIC) can be used to compare models (Tataru and Bataillon 2019). The LRT entails fitting two nested models, where one reduced model is a special case of a more general model. A *P*-value is obtained by assuming that the log of the ratio of the maximum likelihoods of the two models follows a *X^2^* distribution parameterized by the difference in the number of degrees of freedom (i.e. number of estimated parameters) between the two models. A *P*-value below 5% means that the reduced model is rejected in favor of the more parameter-rich model. LRTs are performed comparing the pairs of models indicated in the main text (see Table 1 and 2 for the lists of models). This approach of comparing the DFE parameters between species through LRTs has been applied before (Huber *et al*. 2017; Zhen *et al*. 2018). Simulations demonstrate that the method guards against excessive type I error while retaining substantial power to detect differences across data sets when the *r_i_* parameters are estimated independently for each data set (Tataru and Bataillon 2019). We use the R function *compareModels*() (from: https://github.com/paula-tataru/polyDFE/blob/master/postprocessing.R) to compare pairs of models.

### Subsampling

To assess the contribution of the variation in *s_d_* and *N_e_* to the realized strength of purifying selection (*S_d_*) across great apes, we run a multiple linear regression. We first obtained statistically independent measures of *S_d_* and *s_d_*. Note that *s_d_* = *S_d_* / 2*N_e_*. To do so, we divided the coding genome into two halves drawing random sites without replacement. We use the first half to estimate *S_d_*_,1_ and the second half to estimate *s_d_*_,2_. To estimate *s_d_*_,2_ we divided *S_d_*_,2_ by 2*N_e_*_,1_, where *N_e_*_,1_ is estimated by dividing *θ* (Watterson 1975), estimated using 4-fold synonymous sites coming from the first half of the genome, by the genome-wide mutation rate per site and generation (*μ* = 1.65 x 10^-8^). To obtain an independent estimate of *N_e_* we used 4-fold synonymous sites of the second half of the genome. We repeated this subsampling 100 times. We then run a multiple linear regression in a log-log scale with the R function *lm*(log(*S_d_*_,1_) ∼ log(*s_d_*_,2_) + log(*N_e_*_,2_)). As a sanity check we run the complementary analysis *lm*(log(*S_d_*_,2_) ∼ log(*s_d_*_,1_) + log(*N_e_*_,1_)) obtaining equivalent results (data not shown). The relative importance of *s_d_* and *N_e_* is assessed using standardized regression coefficients (*β*).

### Other Statistical Analyses

All statistical analyses commented above have been performed within the R framework (version 3.4.4) except the DFE estimation and the phylogenetically aware regression. To perform the phylogenetically aware regression between *N_e_* and various summary statistics of the full DFE we use BayesTraitsV3 (Pagel and Meade 2006). We set the method to employ random-walk and maximum likelihood. Significance is assessed by comparing a model where the correlation is a fitted parameter to a model where the correlation is fixed to 0, by means of a likelihood ratio statistic. To perform the bivariate correlations, we use the function *cor.test()* and the Pearson method. To perform the quadratic regression between *N_e_* and *S_d_* (or *s_d_*) we use the R function *lm*(y∼ poly(x, degree = 2)). We then compare the linear regression and polynomial regression with a likelihood ratio test. All figures are generated using the R package “ggplot2”.

### Data Availability Statement

The final list of analyzed positions is available upon request. The authors confirm that all data necessary for confirming the conclusions of the article are present within the article, figures, and tables.

## Results

We compiled polymorphism data from nine great ape populations: bonobos, western chimpanzees, central chimpanzees, Nigeria-Cameroon chimpanzees, eastern chimpanzees, western lowland gorillas, Sumatran orangutans, Bornean orangutans and humans. A cosmopolitan sample of humans (3 African and 6 non-African diploid individuals) was used to represent human genetic diversity. All our population data is retrieved from Prado-Martinez *et al*. (2013). To allow a fair comparison each population is sub-sampled to 8 haploid chromosomes. We investigate more than 9,000 autosomal coding genes that are orthologous one-to-one across great apes, gibbons, and macaques (the two outgroups we use to calculate the unfolded site frequency spectrum, uSFS). See Supplementary Table 1 for the whole list of analyzed genes and the uSFS. The proportion of shared SNPs between species is below 1% but it raises to 10-30% between pairs of chimpanzee populations and between the orangutan species, respectively (Supplementary Table 2). This imperfection of the data makes the chimpanzee populations and orangutan populations statistically non-independent.

The inference of the DFE from the uSFS can be affected by polarization errors (Hernandez *et al*. 2007), GC-biased gene conversion (Bolívar *et al*. 2018), tight linkage (Eyre-Walker and Keightley 2009; Kousathanas and Keightley 2013), ascertainment bias, non-random sampling, population structure and demographic changes (Eyre-Walker *et al*. 2006; Keightley and Eyre-Walker 2007; Boyko *et al*. 2008). To quantify and correct polarization errors, we estimate a polarization error parameter (*ε_anc_*) (Williamson *et al*. 2005; Boyko *et al*. 2008; Glémin *et al*. 2015). To account for additional biases, we compare the observed uSFS for putatively neutral (4-fold degenerate) synonymous mutations to the expected uSFS for neutral mutations under the Wright-Fisher model. To do that, we estimate a series of nuisance parameters (*r_i_*) using the uSFS for 4-fold synonymous and 0-fold non-synonymous mutations (as in Eyre-Walker *et al*. [2006]). If the *r_i_* parameters are significantly different from 1, we jointly estimate the *r_i_* parameters together with the parameters of the DFE, otherwise only the DFE parameters are estimated. The *r_i_* parameters are adjusted for each allele frequency class in the uSFS. This effectively assumes that the neutral and selected uSFS are distorted in an identical way by these factors. Weakly beneficial mutations can also distort the uSFS and by extension our estimates of the deleterious DFE (Tataru *et al*. 2017). We, therefore, also co-estimate the rate and strength of new beneficial mutations together with the *r_i_*, *ε_anc_* and the two parameters defining the deleterious DFE: the shape, *b*, and mean, *S_d_*. Here we assume, like several studies in humans before, that the deleterious DFE is gamma distributed (Boyko *et al*. 2008; Eyre-Walker and Keightley 2009; Galtier 2016; Huber *et al*. 2017; Kim *et al*. 2017). We investigate two beneficial DFEs: an exponential and a more general discrete distribution. Given the very similar results obtained with these two beneficial DFEs in the main text we will only show the results obtained with the exponential distribution. The results obtained assuming a discrete distribution are reported in the Supplementary Table 4.

### Model choice

Before comparing the DFE between species we used a variety of models to fit our data at the species level. Differences among models include the joint estimation of DFE parameters and extra nuisance parameters that can correct for departures from a constant population size Wright-Fisher model, the proportion of beneficial mutations, and potential polarization errors. Table 1 shows the different DFE models and parameters used in this work.

We find that the uSFS of all great apes show significant departures from the constant population size Wright-Fisher model (*r_i_* ≠ 1 as judged by a likelihood-ratio test, LRT) (Supplementary Table 3, column M0 vs M1). These nuisance parameters capture all evolutionary processes that equally affect synonymous and non-synonymous mutations, such as demography, population structure, sampling bias, and linked selection. The performance of these nuisance parameters under demographic histories relevant for great apes is investigated using simulations presented in the Supplementary Material. In general, the inference quality is very high with simulated data, particularly when some aspects of the DFE are jointly estimated across species (see below).

We find that the models including *ε_anc_* (M2 and M4) do not fit the data significantly better than the models assuming *ε_anc_* = 0 (M1 and M3) (Supplementary Table 3, column M1 vs M2 and M3 vs M4), except in eastern chimpanzees. This indicates that our polarization strategy is effective or that a substantial fraction of polarization errors is accounted for by the *r_i_* parameters (as argued before by Galtier [2016]). We do not find strong evidence for beneficial amino acid mutations segregating in our samples (Supplementary Table 3, column M1 vs M3 and M2 vs M4). Consistent with these findings, the most flexible model (M2) is never the preferred model, as judged by our LRTs (Supplementary Table 3). M3 is instead the preferred model across all species. This model assumes only deleterious mutations and no polarization errors but includes the distortion parameters (*r_i_*). The modest contribution of beneficial mutations is corroborated by the goodness-of-fit analyses (Figure 1). The results comparing the exponential and discrete beneficial DFE can be consulted in Supplementary Table 4. Both distributions fit the data equally well. Larger samples sizes will be needed to assess which beneficial distribution is more realistic. Hereafter, we will refer to M3 as the purely deleterious model and M2 as the flexible model.

**Figure 1.**
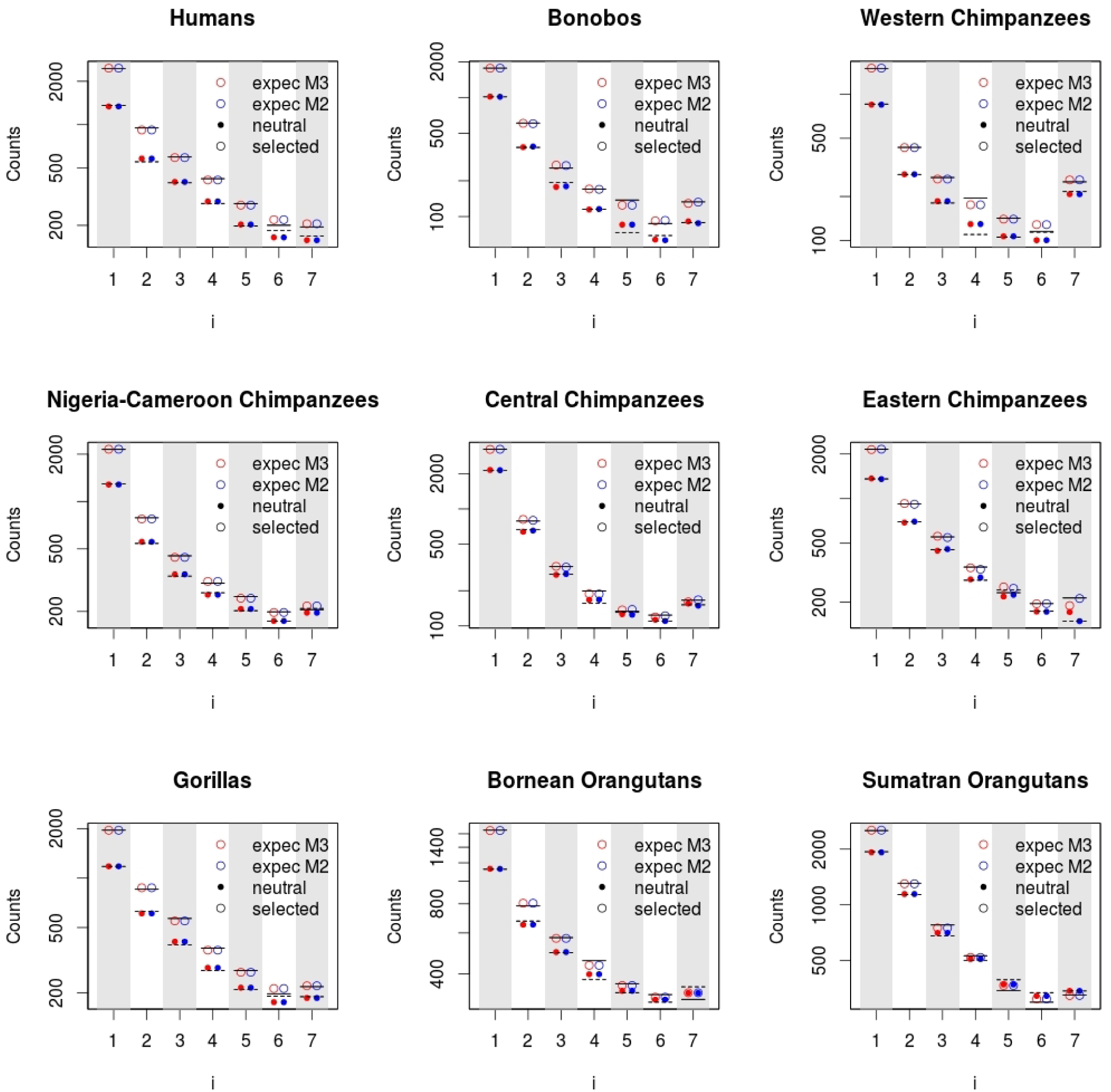
Observed and expected uSFS. The solid lines show the observed number of 0-fold non-synonymous changes and the dashed lines show the observed number of 4-fold synonymous changes. The dots and circles represent the expected counts of neutral and selected sites under the M2 and M3 models (shown as blue and red, respectively). The y-axis is on log scale.

### Assessing the stability/invariance of the DFE across great apes

Next, we sought to investigate whether the full DFE varies or not across great apes. We compare two models: one where only the shape parameter (*b*) of the deleterious DFE is shared across species and where the mean, *S_d_*, can vary independently as expected for populations with different effective sizes, the other where all the parameters, including *b*, vary independently across species (this is equivalent to model 3 in Table 1). Note that, in a few cases, shape estimates of the deleterious DFE can be very sensitive to the model assumptions and in particular whether the model includes beneficial mutations (M1 and M2) or not (M3 and M4). For example, in bonobos the estimated shape parameter is *b* = 0.09 under the purely deleterious model, but it increases to *b* = 0.22 under the flexible model, which includes beneficial mutations (Supplementary Table 4). This is because under the flexible model a substantial proportion of new mutations are inferred as slightly beneficial or nearly neutral (*p_b_* ∼ 14-17% and average *S_b_* ∼ 0.65-0.79). In other words, in bonobos the shape of the deleterious DFE is sensitive to the presence/absence of beneficial mutations in the model. Hence, to accommodate these cases we also include the flexible model where both the beneficial DFE parameters and *ε_anc_* are jointly estimated.

Table 2 shows the details of the set of constrained models used to test DFE invariance. In all these models the distortion parameters (*r_i_*) are estimated independently to each species as recommended by Tataru and Bataillon (2019). Figure 2 shows that the density distributions for the shape of the deleterious DFE largely overlap between species. We find that the model estimating the shape of the deleterious DFE independently in each species does not explain the data significantly better than the model where the shape parameter is shared across species (shared *b* = 0.1588, bootstrap confidence interval 95% = [0.1344, 0.1736]) (M3S vs M3I: LRT *P*-value = 0.77). Although visually it seems that bonobos show a shift toward lower shape parameters, this result is not significant (M3 independent *b* = 0.0938 vs M3 shared *b* = 0.1588: LRT *P*-value = 0.07). The inferred shape of the deleterious DFE increases very slightly when beneficial mutations are also modeled (shared *b* = 0.1842 [0.1697, 0.2184]) (M2S vs M2I: LRT *P*-value = 0.97) (Supplementary Table 5). Moreover, we find that the model where the shape of the deleterious DFE is shared across species but the beneficial DFE is estimated independently for each species (M2S) does not fit the data significantly better than the equivalent model without beneficial mutations (M3S) (M3S vs M2S: LRT *P*-value = 0.99).

**Figure 2.**
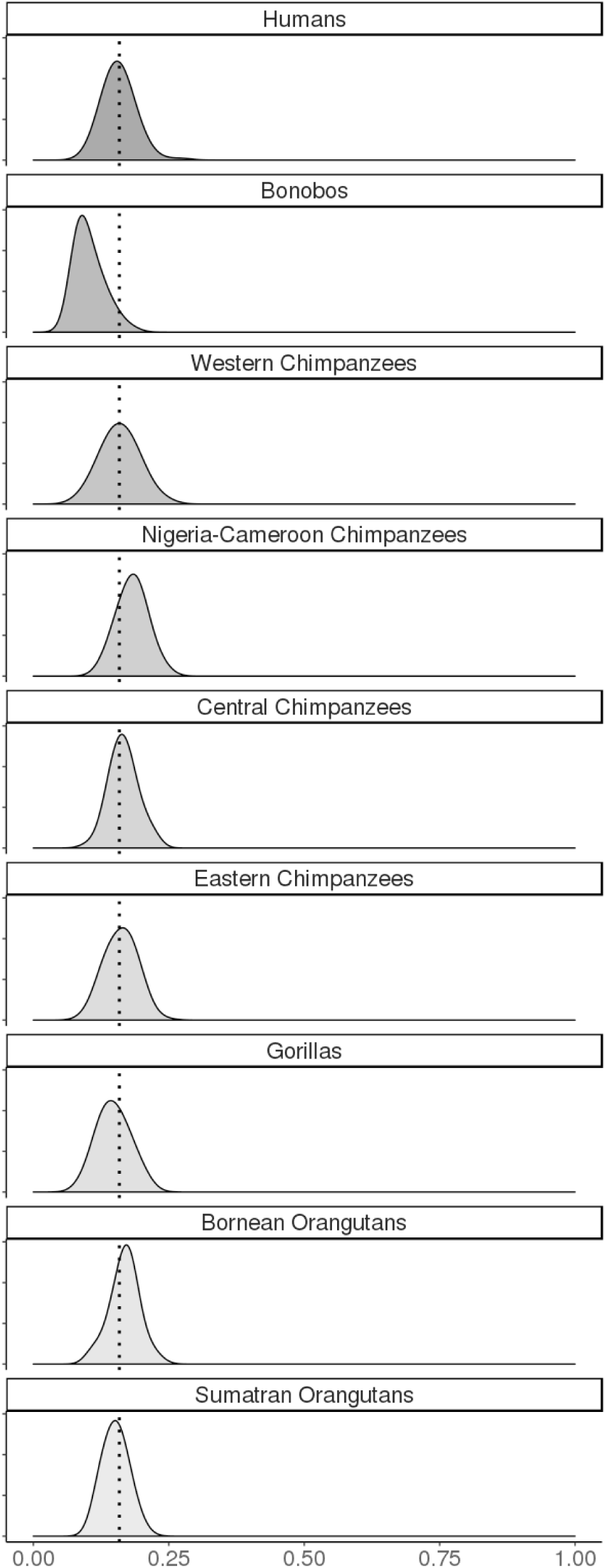
Sampling distribution of shape parameter estimates (*b*) for the DFE estimated in each species. Sampling distributions are obtained by 100 bootstrap replicates for each species and estimating *b* under the purely deleterious model (M3I). The dotted black line (*b* = 0.1588) indicates the mean of the shared shape parameter under the most likely model (M3S). See Table 2 for a description of the models.

**Table 2.** List of estimated shared and independent parameters for each model. Models 2I and 3I are equivalent to models 2 and 3 from Table 1, respectively.

| Model | $S_d$ | $b$ | $S_b$ | $p_b$ | $r_i$ | $\epsilon_{anc}$ | $\theta$ |
| --- | --- | --- | --- | --- | --- | --- | --- |
| **2I** | independent | independent | independent | independent | independent | independent | independent |
| **2S** | independent | shared | independent | independent | independent | independent | independent |
| **3I** | independent | independent | 0 | 0 | independent | 0 | independent |
| **3S** | independent | shared | 0 | 0 | independent | 0 | independent |
| **3SS** | shared | shared | 0 | 0 | independent | 0 | independent |

Population genetics theory predicts that natural selection will be weaker in small populations due to random genetic drift (Ohta 1992). If we assume that the shape of the deleterious DFE is constant across great apes, we find a strong positive correlation (Pearson’s correlation coefficient *r* = 0.91 [0.88, 0.93]) (Figure 3 A) between the population scaled mean effect size of deleterious mutations (|*S_d_*| = |2*N_e_s_d_*|) and our estimate of the effective population size (*N_e_* = θ_S_ / (4*μ*), where *θ_S_* is synonymous diversity and *μ* the mutation rate per site and generation, *μ =* 1.65 x 10^-8^) (Ségurel *et al*. 2014). This correlation remains significant after accounting for species phylogenetic dependence (BayesTrait V3 *P*-value < 0.01). Forward simulations confirm that *S_d_* and *N_e_* are significantly correlated as expected by the Nearly Neutral Theory (Supplementary Analyses). We also find that the mean selection coefficient of deleterious mutations, *s_d_* (calculated dividing *S_d_* by *2N_e_*), is positively correlated to *N_e_* (*r* = 0.76 [0.67, 0.85], BayesTrait V3 *P*-value < 0.01). We checked using simulations that our method of inference does not introduce any co-variation between *s_d_* and *N_e_* (Supplementary Analyses). We then assessed the relative contribution of the variation in *N_e_* and *s_d_* to the differences in *S_d_*. To do so, we use a multiple linear regression. *S_d_* estimates are subject to considerable uncertainty and statistical noise and this can introduce a spurious correlation between *S_d_* and *s_d_* estimates. Hence, we first obtained *S_d_* and *s_d_* estimates that are statistically independent by splitting the coding genome in two (as in James *et al*. 2016; Castellano *et al*. 2016, 2018a) (see Material and Methods: Subsampling). One half is used to estimate *S_d_* and the other half to estimate *s_d_*. We run one hundred replicates to compute the empirical confidence intervals. We find that *N_e_* and *s_d_* are both significantly correlated with *S_d_* in our multiple linear regression (Table 3). The standardized regression coefficients (*β*) suggest that direct variation in *N_e_* and *s_d_* are quantitatively of very similar magnitude (*β*(*N_e_*) = 0.43 [0.22, 0.85] and *β*(*s_d_*) = 0.69 [0.47, 1.07]). Although our predictor variables (*N_e_* and *s_d_*) are correlated, their variance inflation factors are small (VIF ∼ 1.1) suggesting that multicollinearity is a minor issue.

**Figure 3.**
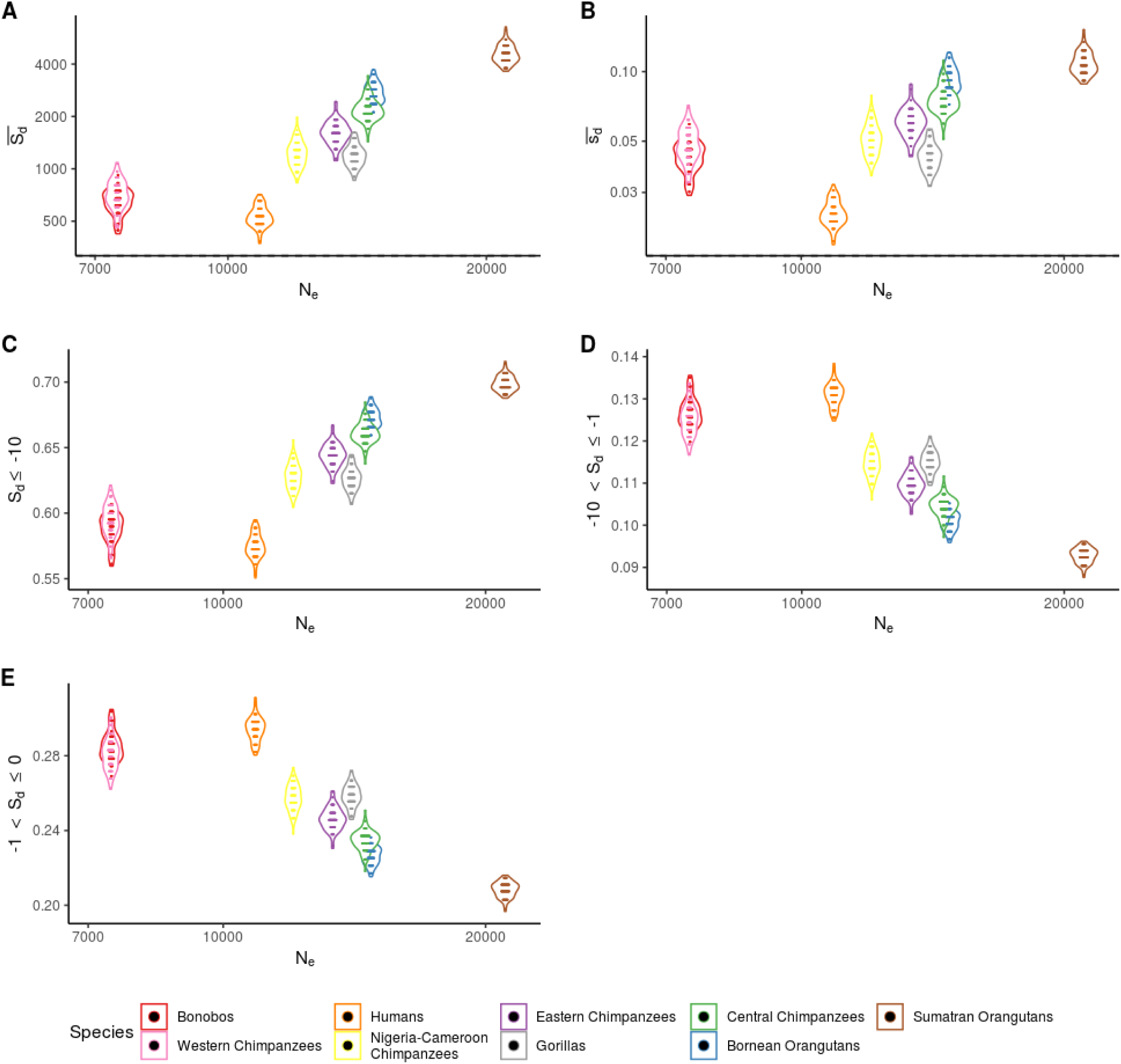
Relationship between *N_e_* and *S_d_* on log scale (A), *S_d_* on log scale (B), the proportion of strongly deleterious mutations (C), the proportion of weakly deleterious mutations (D) and the proportion of effectively neutral mutations (E). Note that *N_e_* (x-axis) is visualized on log scale. Each violin plot represents 100 bootstrap replicates under the most likely model (M3S). See Table 2 for a description of the models.

**Table 3.** Regression analysis. A’, B’ and C’ refer to the models with only humans, bonobos, western and central chimpanzees to make a direct comparison with the simulated data (see Supplementary Analyses).

| Model | Slope | Adjusted R<sup>2</sup> | P-value |
| --- | --- | --- | --- |
| $A: \log(S_{d,1}) \sim \log(N_{e,2})$ | 1.90 (1.64, 2.16) | 0.70 | < 0.001 |
| $B: \log(s_{d,2}) \sim \log(N_{e,2})$ | 0.88 (0.63, 1.13) | 0.34 | < 0.001 |
| $C: \log(S_{d,1}) \sim \log(N_{e,2}) + \log(s_{d,2})$ | $N_e$: 1.35 (0.98, 1.50)<br>$s_d$: 0.63 (0.46, 0.80) | 0.81 | < 0.001 |
| $A'$ | 1.42 (1.29, 1.58) | 0.40 | < 0.001 |
| $B'$ | 0.52 (0.01, 1.02) | 0.08 | < 0.05 |
| $C'$ | $N_e$: 1.02 (0.57, 1.48)<br>$s_d$: 0.68 (0.36, 0.85) | 0.63 | < 0.001 |

Of note, although a linear regression explains most of the variance in *S_d_* (*R^2^* = 0.83 [0.77, 0.86]), the relationship seems to reach a plateau for the less populous great apes. In fact, we find that a curvilinear function (quadratic regression) fits the data significantly better (*R^2^* = 0.90 [0.80, 0.95]) than a linear regression (LRT linear vs quadratic regression *P*-value < 0.01). This result is not observed in the simulated data sets (Supplementary Analyses) and it suggests that other forces beyond genetic drift can modulate the strength of purifying selection in the smallest great ape populations (see Discussion).

One can argue that our sample sizes are too small to estimate with precision *S_d_* and that the findings commented above are merely driven by noise or by some particular demography biasing the estimates of the DFE parameters inferred using polyDFE. However, estimates of the proportion of mutations in a given *S_d_* range from the DFE are often less noisy than mean *S_d_* estimates (Keightley and Eyre-Walker 2007; Eyre-Walker and Keightley 2009). We demonstrate with our simulations that when the shape parameter is co-estimated across species, not only these proportions can be reliably estimated using a small sample size, but also *S_d_* (Supplementary Analyses). We find that the fraction of strongly deleterious mutations (*S_d_* ≤ −10) is positively correlated to *N_e_* (Figure 3 C, Table 4). There is a negative correlation between the fraction of effectively neutral mutations (−1 < *S_d_* ≤ 0) and *N_e_* (Figure 3 E, Table 4) as predicted by the Nearly Neutral Theory (Ohta 1992). Interestingly, there is a negative, but very shallow, correlation between the fraction of weakly deleterious mutations (−10 < *S_d_* ≤ −1) and *N_e_* (Figure 3 D, Table 4). This is consistent with a recent work that found that the impact of linked selection (mainly background selection) on genetic diversity is very similar along the genomes of great apes (Castellano *et al*. 2018b).

**Table 4.**
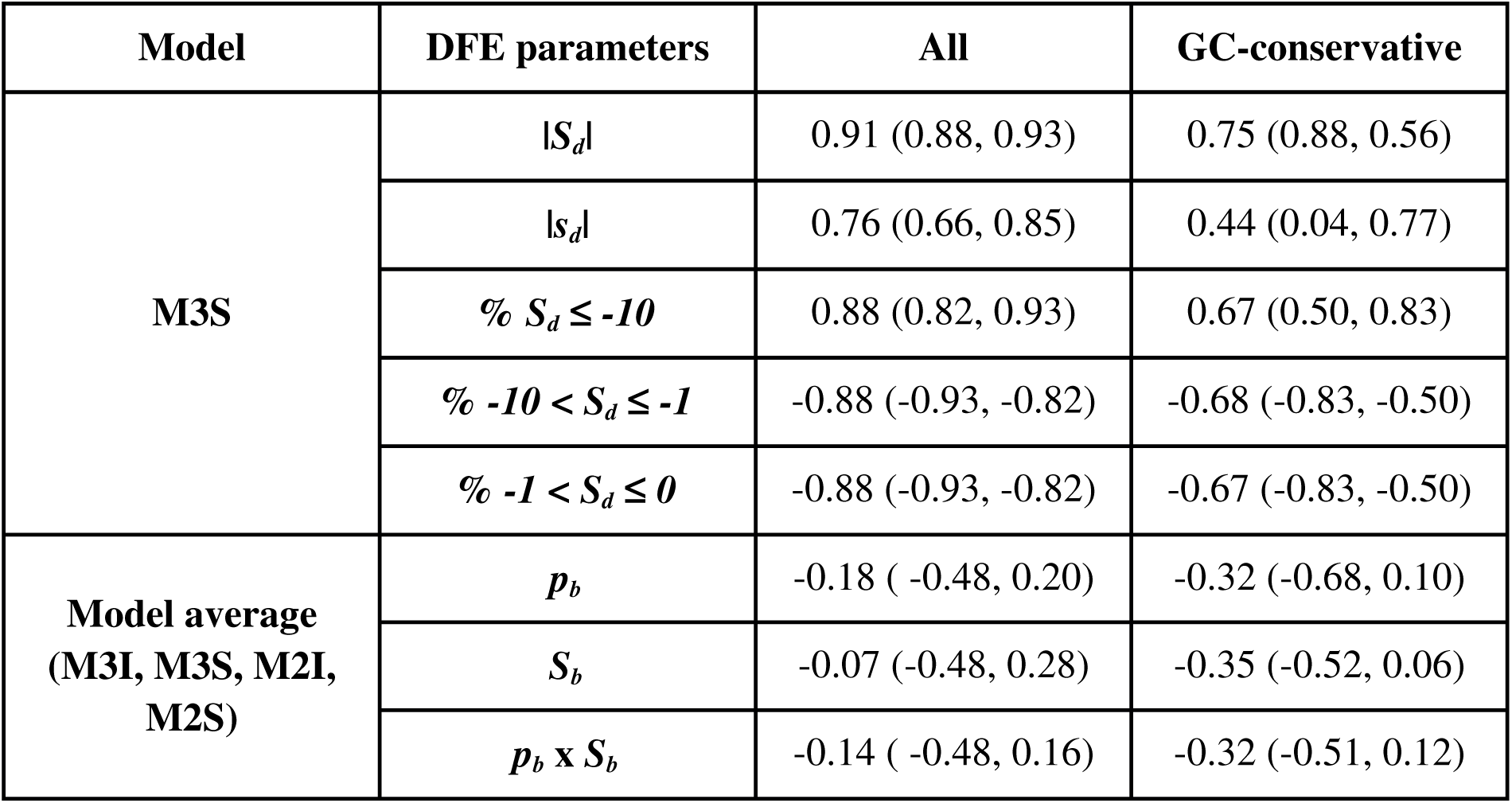
Pearson correlation coefficient between the parameters describing the full DFE and *N_e_* for all amino acid changing mutations and GC-conservative mutations. In parentheses the 95% confidence interval based on 100 bootstrap replicates. All these correlations have been cross-validated using a phylogenetically aware regression analysis.

To further assess whether there is systematic variation in *S_d_* across great apes, we compare a purely deleterious model where both *b* and *S_d_* are shared across all species (M3SS) against a model where only *b* is shared between species (M3S). In both models, the *r_i_* and the population scaled mutation rate parameters are estimated independently for each species. The model with variation in the mean *S_d_* across great apes fits the data significantly better than the model where the mean *S_d_* is the same across species (M3S vs M3SS: LRT *P*-value = 5.9e^-40^) (Supplementary Table 5 E). This result strongly supports the fact that there is systematic variation in the strength of purifying selection across the great apes.

The selective effect, *S_b_*, of new beneficial mutations might also depend on *N_e_*. For instance, some authors have reported that species with a larger *N_e_* tend to have higher rates of beneficial substitutions (Strasburg *et al*. 2011; Gossmann *et al*. 2012). This is because large populations will have to wait less time for the appearance of new beneficial mutations. Once the beneficial mutation has appeared, natural selection will be more effective in populations with large *N_e_*. In contrast, other studies have reported that proxies for *N_e_* and the rate of beneficial substitutions are poorly correlated (Galtier 2016). This is expected if small *N_e_* populations have higher rates of new beneficial mutations, *p_b_*, because they are further away from their fitness optimum due to the fixation of slightly deleterious mutations and/or the higher genetic load (segregation of deleterious mutations) (Hartl and Taubes 1996; Poon and Otto 2000).

To test whether our data support any of those opposing views we use a model averaging approach to estimate the effect and proportion of new beneficial mutations (Tataru and Bataillon 2019). This allows us to factor in the fact that, as measured via LRT, there is only very weak (statistically non-significant) evidence for the presence of beneficial mutations in the polymorphism data. We consider a set of four competing models (M3I, M3S, M2I and M2S) and weight them by their AIC. Supplementary Figure 1 A shows the weight of each model per species. This approach has been applied before in the context of detection of adaptive molecular evolution (Kjeldsen *et al*. 2012; Rousselle *et al*. 2019). We find a non-significant negative correlation between the model-averaged rate of new beneficial mutations, *p_b_*, and *N_e_* (Table 4, Supplementary Figure 2 A). There is no correlation between the model-averaged *S_b_* and *N_e_* (Table 4, Supplementary Figure 2 B) and the expected rate of beneficial non-synonymous substitutions (the product of rate of new beneficial mutations, *p_b_*, and the mean strength of positive selection, *S_b_*) is also uncorrelated to *N_e_* (Table 4, Supplementary Figure 2 C). Model-averaged estimates of the proportion of beneficial mutations, *p_b_*, suggest that beneficial mutations remain very rare (Supplementary Figure 3) and *S_b_* estimates are always small (*S_b_* << 1) and similar across great apes (Supplementary Figure 4). Nevertheless, it is also worth mentioning that one of the two smallest populations of great apes, bonobos, show a substantial proportion (1-2%) of new beneficial mutations (Table 5; Supplementary Figure 3 and 8). Larger sample sizes will be required to quantify this interesting class of effectively neutral beneficial mutations precisely.

**Table 5.** *p_b_* and *S_b_* mean and 95% empirical confidence interval across 100 bootstrap replicates. *p_b_* and *S_b_* estimated weighting models M2I, M3I, M2S and M3S for each species. Species ordered by mean *p_b_*.

| | $p_b$ | $S_b$ |
| --- | --- | --- |
| **Bonobos** | 1.7e-2 (7.3e-5, 1.2e-1) | 0.0760 (0.0008, 27.8058) |
| **Central Chimpanzees** | 9.0e-3 (1.7e-5, 1.2e-1) | 0.0673 (0.0023, 3.4063) |
| **Gorillas** | 1.3e-3 (7.1e-6, 3.0e-2) | 0.0091 (0.0008, 0.2291) |
| **Eastern Chimpanzees** | 4.8e-4 (1.6e-5, 8.7e-2) | 0.1830 (0.0102, 8.3537) |
| **Sumatran Orangutans** | 7.2e-5 (2.2e-6, 6.8e-3) | 0.0080 (0.0010, 0.4206) |
| **Bornean Orangutans** | 7.0e-5 (3.1e-6, 2.3e-2) | 0.0087 (0.0010, 0.1481) |
| **Western Chimpanzees** | 2.9e-5 (3.2e-6, 1.0e-2) | 0.0081 (0.0007, 0.0911) |
| **Humans** | 2.3e-5 (2.2e-6, 7.6e-3) | 0.0064 (0.0007, 0.1084) |
| **Nigeria-Cameroon Chimpanzees** | 1.6e-5 (3.0e-6, 2.6e-3) | 0.0104 (0.0017, 0.1382) |

### Accounting for gBGC

GC-biased gene conversion (gBGC) can distort the uSFS of synonymous and non-synonymous mutations differently due to differences in their GC-content (Bolívar *et al*. 2018). Furthermore, larger populations are expected to be more affected by gBGC due to higher effective rates of recombination and gene conversion. Thus, the degree of bias will also scale with *N_e_*. To check that our findings are not merely driven by differences in the intensity of gBGC, we repeated our analysis using GC-conservative mutations (A<->T and C<->G), which are unaffected by gBGC. Supplementary Table 6 shows the estimated parameters of the DFE for GC-conservative mutations for each species and model. We also find that the model where a single shape *b* is shared across species is preferred over the model with a separate shape parameter for each species (shared *b* = 0.1694 [0.1123, 0.2176]) (M3S vs M3I: LRT *P*-value = 0.63) (Supplementary Figure 5). This is also true when beneficial mutations are included (shared *b* = 0.2789 [0.2014, 0.9997]) (M2S vs M2I: LRT *P*-value = 0.70). We also find evidence of systematic variation in the strength of negative selection for GC-conservative mutations across species. A model where the shape is shared but the mean *S_d_* is estimated independently is preferred over a model where both the shape and the mean *S_d_* are shared (M3S vs M3SS: LRT *P*-value = 4.4e^-6^).

We confirm that *s_d_* co-varies with *N_e_* (Table 4) and that the strength of purifying selection in bonobos and western chimpanzees is higher than expected given their *N_e_* (Supplementary Figure 6). The correlation between *N_e_* and the summary statistics of the DFE also persist when considering only non-synonymous GC-conservative mutations (Table 4; Supplementary Figure 6-7; Supplementary Table 7). The goodness-of-fit analysis for GC-conservative mutations is presented in Supplementary Figure 9.

## Discussion

In this work, we have inferred and compared the full distribution of fitness effects (DFE) of new heterozygous mutations between humans and their closest living relatives. To estimate the DFE we used only allele frequency distributions for 4-fold synonymous and 0-fold non-synonymous changes. We found that the shape of the deleterious DFE is remarkably constant across great apes, *b* = 0.16 (0.13, 0.17). This result is robust to gBGC, polarization errors and proportions and effect of new beneficial mutations. Our estimate of the shape parameter in humans is consistent with older reports (*b* = 0.18-0.20) (Eyre-Walker *et al*. 2006; Keightley and Eyre-Walker 2007; Boyko *et al*. 2008) and recent estimates where thousands of human chromosomes are re-analyzed under a complex human demography (*b* = 0.17-0.21) (Kim *et al*. 2017). While our analysis did not infer specific demographic parameters, co-estimating nuisance parameters for the different uSFS classes does capture the effects of demography and allows us to recover the underlying DFE. We confirmed the robustness of our approach by running forward simulations with demographic histories relevant for great apes (Supplementary Analyses).

If we assume that within the great apes, not only recombination rate, mutation rate, gene density or gene expression levels are conserved but also protein function and regulatory and metabolic networks are equivalent, then it may be possible to explain the differences in the DFE between species by differences in their effective population size. Hence, here we ask: Does *N_e_* affect the DFE all else being equal? We found evidence for systematic variation in the strength of negative selection (*S_d_* = 2*N_e_s_d_*) across great apes. The correlation between our estimate of the species effective population size (*N_e_*), based on the current levels of diversity at 4-fold synonymous sites (*θ_S_*), and the estimated strength of negative selection (*S_d_*) is very strong for all mutations and GC-conservative mutations (explaining ∼80% and ∼60% of the variance in the strength of purifying selection, respectively). This result is consistent with the Nearly Neutral Theory (Ohta 1992) assuming that the deleterious DFE (in *s_d_* units) is constant over evolutionary time. This constancy is expected under a model where protein function is the main driver of fitness effects of mutations, and where protein function hardly changes between species (at the evolutionary time scale spanning great apes). However, we also found evidence that suggests that *s_d_* has not remained constant across great apes. We expect that *S_d_* scales proportionally with *N_e_*. Surprisingly, we found more pronounced differences in *S_d_* than expected given our estimates of the effective population size based on current levels of synonymous diversity. In other words, we find that the mean absolute effect size of deleterious mutations (*s_d_*) is also correlated to *N_e_*. Using data simulated with a constant *s_d_*, we show that our estimation procedure does not drive the co-variation between *N_e_* and *s_d_* estimates. Interestingly, this result is consistent with positive epistasis. Under positive epistasis *s_d_* will increase as fitness (and probably *N_e_*) decreases (Silander *et al*. 2007). This is because new deleterious mutations will be less detrimental in a genetic background that already contains deleterious variants than in a genetic background free of deleterious mutations. Population genetics theory predicts that in small populations drift can overwhelm selection. Slightly deleterious mutations may thus reach higher frequencies and even reach fixation causing fitness to decline (Kimura *et al*. 1963; Bataillon and Kirkpatrick 2000). This means that new deleterious mutations will have a higher chance of interacting with other pre-existing deleterious variants in a small population than in a large population. The prevalence of epistasis is thus expected to increase when *N_e_* decreases and this is reflected in our estimates of the DFE. We propose that the variation in the strength of purifying selection across great apes is doubly affected by *N_e_*. First, for a given selection coefficient, *N_e_* determines the efficiency of purifying selection (Ohta 1992) and second, *N_e_* determines the amount of potential epistatic interactions occurring in a given individual which in turn will affect the magnitude of the effect of new deleterious mutations (Poon and Otto 2000). This is an exciting result that deserves further theoretical exploration and empirical validation.

Moreover, for additive mutations, the effect of deleterious mutations should decrease with decreasing *N_e_*. Surprisingly, we find that in bonobos and western chimpanzees, the two smallest great ape populations, the mean effect size of deleterious mutations increases. We do not see this overestimation of *S_d_* in our forward simulations of the bonobo and western chimpanzee demographic histories. Note that in our simulations all mutations are codominant. We hypothesize that the efficient purging of strongly deleterious recessive variants in bonobos and western chimpanzees might explain this result (Barrett and Charlesworth 1991; Glémin 2003). Bonobos and eastern lowland gorillas (not analyzed in this study) show an excess of inbreeding compared to the other great apes, suggesting small population sizes or a fragmented population (Prado-Martinez *et al*. 2013). Similarly, the western chimpanzees are thought to have spread from a very small ancestral population (Prado-Martinez *et al*. 2013; de Manuel *et al*. 2016) and show, as expected by theory, a higher proportion of putatively deleterious variants compared to central chimpanzees (Han *et al*. 2019). Recent work in *Ibex* suggests that bottlenecks can indeed favor the purging of strongly deleterious recessive mutations while allowing the accumulation of weakly deleterious additive mutations (Grossen *et al*. 2019). Note, however, that our estimates assume that all mutations segregating and affecting fitness are codominant (*h* = 0.5). Whether the estimation of DFE parameters is robust to variation in the joint effects of the dominance of mutations, inbreeding and demography remain an open question.

Regarding the beneficial portion of the DFE, we do not find any statistically significant contribution of beneficial mutations to the uSFS counts. Thus, we are unable to support either an increase in the strength of positive selection (*S_b_*) with *N_e_* (Nam *et al*. 2017), or an increase in the expected rate of beneficial substitutions (*p_b_* x *S_b_*) with *N_e_* (Eyre-Walker 2006). Note, however, that in this work we do not use substitution data, only polymorphisms. Rare and strongly beneficial mutations will fix quickly and contribute relatively more to divergence counts than to uSFS counts (Eyre-Walker and Keightley 2007). Hence, our results are still compatible with the view that a sizeable amount of divergence at the amino acid level is driven by relatively rare but strongly beneficial mutations in the great apes (Nam et al. 2017). Our choice was to avoid using divergence data to estimate the full DFE. The reason for doing so is that fixed mutations may introduce biases in the estimation of the full DFE if ancient fluctuations in the effective population size are not properly modelled (Tataru *et al*. 2017; Rousselle *et al*. 2018; Zhen *et al*. 2018). A second explanation to the apparent lack of beneficial mutations is our modest sample size (n = 8 haploid chromosomes per population). Thus, although most adaptive substitutions seem to be weakly beneficial in humans (Uricchio *et al*. 2019), we might need very large sample sizes to quantify accurately the DFE of new beneficial mutations because new beneficial mutations are still very rare relative to deleterious ones.

Using a model averaging framework where the different competing models are weighted by their AIC, we find that between 1-2% of new mutations are mildly beneficial in bonobos. However, the estimated population scaled effect size of beneficial mutations is below one in all great apes. Small populations tend to be further away from their fitness optimum due to a higher genetic load and/or fixation of slightly deleterious mutations. As a consequence, a new mutation has a higher probability of being beneficial in a small than in a large population (Hartl and Taubes 1996; Poon and Otto 2000; Silander *et al*. 2007). The interactions within and among genes will allow new mutations to compensate/restore for the fitness effects of other, fixed or polymorphic, slightly deleterious mutations. Hence, there is no need for very unlikely back-mutations to restore the fitness losses incurred by previous mutations (Charlesworth and Eyre-Walker 2007). An interesting implication of such mode of evolution is that rates of adaptive substitutions may not be driven only by external conditions (such as viruses, see Enard *et al*. 2016; Castellano *et al*. 2019; Uricchio *et al*. 2019) but also by the amount of deleterious mutations already present in the genome as this mutation load conditions the current level of adaptation in a population. This mechanism is not often invoked to explain Darwinian adaptation (due to environmental changes), yet a small pool of compensatory mutations will contribute to the amino acid differences between species in the long-term (Hartl and Taubes 1996). The induced epistasis imply that mutations are only conditionally beneficial or deleterious and that the DFE and *N_e_* might not be independent as commonly assumed.

Finally, we discuss some limitations of our study. We have assumed the same mutation rate in all populations to estimate *N_e_* based on current levels of synonymous diversity, *θ_S_*. Synonymous diversity has been used repeatedly in several related studies as a proxy for *N_e_* (Gossmann *et al*. 2010; Strasburg *et al*. 2011; Phifer-Rixey *et al*. 2012; Galtier 2016). This might be a problem because *θ_S_* is jointly influenced by variation in *N_e_* and also by the mutation rate. In this work we have assumed the same mutation rate per site and generation across all great apes. However, there is evidence that both the generation time and the mutation rate per year vary across great apes (Amster and Sella 2016; Jónsson *et al*. 2017; Thomas *et al*. 2018; Besenbacher *et al*. 2019). We checked how more realistic estimates of mutation rate per generation could explain our results. With data retrieved from Besenbacher *et al*. (2019) we find that the mutation rate per generation is ∼23% higher in chimpanzees, bonobos and orangutans than in humans. Gorillas and humans have a very similar mutation rate per generation despite having the shortest and longest generation times across great apes, respectively. We show that our results, including the correlation between *s_d_* and *N_e_*, remain robust to the variation in the mutation rate per generation across great apes (Supplementary Figure 10; Supplementary Table 8). We emphasize that estimates of mutation rate per generation are prone to much uncertainty due to our limited knowledge about generation times in nature. Second, synonymous diversity is a poor proxy for *N_e_* if there is widespread selection on codon usage or other kinds of selection on synonymous changes due to, for example, the regulation of splicing or gene expression. However, this is unlikely to affect some great apes more than others or to affect more synonymous than non-synonymous changes. Selection on codon usage is a weak force and great apes are in general not highly populous species. The robustness of our results to gBGC, which is typically correlated to GC-content and codon usage bias, does not suggest that this is a major issue with this analysis. Furthermore, the fact that all our results have been replicated using GC-conservative mutations indicates that gBGC is not likely affecting our conclusions. Third, we cannot explain why we do not observe signals of mildly beneficial mutations in western chimpanzees, a population with a level of genetic diversity equivalent to that found in bonobos. Other factors beyond the genome-wide amount of DNA diversity, such as the rate of change of the environment (Lourenço *et al*. 2013), inbreeding or population structure, might be triggering the emergence of those weakly beneficial mutations specifically in bonobos. Fourth, fluctuations in the effective population size (along time or the genome) might affect neutral and selected mutations differently (Gordo and Dionisio 2005; Brandvain and Wright 2016; Castellano *et al*. 2018c; Torres *et al*. 2019). In other words, the *N_e_* that better describes the population dynamics of neutral mutations might be different from the *N_e_* that better describes the population dynamics of weakly selected mutations or strongly selected mutations (Messer and Petrov 2013; Rousselle *et al*. 2018). These de-couplings in the *N_e_* of neutral and selected mutations can affect our results and interpretations. However, the high inference accuracy of polyDFE under background selection and the complex demographic histories of great apes should make our inference robust to such non-equilibrium dynamics.

## Conclusions

We have made the first comparison of the full distribution of fitness effects of new amino acid mutations across great apes. By comparing the fit of a series of nested models to polymorphism data we have identified which aspects of the DFE are shared between humans and their closest living relatives. Our analysis shows that the shape of the deleterious DFE is shared across these species. Consistent with the Nearly Neutral Theory we have found that the average population scaled effect size of new deleterious mutations (*S_d_*) is strongly correlated to our estimate of *N_e_*. Interestingly, there is also co-variation between the average effect size of new deleterious mutations (*s_d_*) and *N_e_* suggesting a role for positive epistasis. We also find that the two smallest great ape populations, western chimpanzees and bonobos, show a comparatively larger strength of purifying selection, which is compatible with the efficient purging of deleterious recessive variants in small populations. The LRT does not favor a richer DFE model including beneficial mutations over models considering only deleterious mutations. However, when we use a model averaging approach, we estimate a small proportion of mildly beneficial mutations only in bonobos. This finding is consistent with compensatory epistasis, which predicts a larger rate of beneficial mutations in small populations. This work invites further investigation of the relationship between epistasis and *N_e_*. Our study demonstrates the simple but perhaps underappreciated fact, that the effect of mutations is dynamic, even between closely related species, and may depend on the genetic background on which they arise.

## Acknowledgments

We thank Martin Lascoux, Donate Weghorn and Miguel Rodriguez for helpful discussion. We further thank Alex Cagan for sharing the set of sites with a unique mapping to the human genome with at least 5-fold coverage in all individuals per species. This work was supported by the Danish Council For Independent Research (grant number 4181-00358).

## References

Amster G., and G. Sella, 2016 Life history effects on the molecular clock of autosomes and sex chromosomes. Proceedings of the National Academy of Sciences 113: 1588–1593.

Barrett S. C. H., and D. Charlesworth, 1991 Effects of a change in the level of inbreeding on the genetic load. Nature 352: 522.

Barton H. J., and K. Zeng, 2018 New Methods for Inferring the Distribution of Fitness Effects for INDELs and SNPs. Mol. Biol. Evol. 35: 1536–1546.

Bataillon T., and M. Kirkpatrick, 2000 Inbreeding depression due to mildly deleterious mutations in finite populations: size does matter. Genet. Res..

Bataillon T., J. Duan, C. Hvilsom, X. Jin, Y. Li, et al., 2015 Inference of Purifying and Positive Selection in Three Subspecies of Chimpanzees (Pan troglodytes) from Exome Sequencing. Genome Biol. Evol. 7: 1122–1132.

Besenbacher S., C. Hvilsom, T. Marques-Bonet, T. Mailund, and M. H. Schierup, 2019 Direct estimation of mutations in great apes reconciles phylogenetic dating. Nat Ecol Evol 3: 286–292.

Bolívar P., C. F. Mugal, M. Rossi, A. Nater, M. Wang, et al., 2018 Biased Inference of Selection Due to GC-Biased Gene Conversion and the Rate of Protein Evolution in Flycatchers When Accounting for It. Mol. Biol. Evol. 35: 2475–2486.

Boyko A. R., S. H. Williamson, A. R. Indap, J. D. Degenhardt, R. D. Hernandez, et al., 2008 Assessing the evolutionary impact of amino acid mutations in the human genome. PLoS Genet. 4: e1000083.

Brandvain Y., and S. I. Wright, 2016 The limits of natural selection in a nonequilibrium world. Trends Genet.

Cagan A., C. Theunert, H. Laayouni, G. Santpere, M. Pybus, et al., 2016 Natural Selection in the Great Apes. Mol. Biol. Evol. 33: 3268–3283.

Castellano D., M. Coronado-Zamora, J. L. Campos, A. Barbadilla, and A. Eyre-Walker, 2016 Adaptive Evolution Is Substantially Impeded by Hill–Robertson Interference in Drosophila. Mol. Biol. Evol. 33: 442–455.

Castellano D., J. James, and A. Eyre-Walker, 2018a Nearly Neutral Evolution across the Drosophila melanogaster Genome. Mol. Biol. Evol. https://doi.org/10.1093/molbev/msy164

Castellano D., A. Eyre-Walker, and K. Munch, 2018b Impact of mutation rate and selection at linked sites on fine-scale DNA variation across the homininae genome. bioRxiv 452201.

Castellano D., J. James, and A. Eyre-Walker, 2018c Nearly Neutral Evolution across the Drosophila melanogaster Genome. Mol. Biol. Evol. 35: 2685–2694.

Castellano D., L. H. Uricchio, K. Munch, and D. Enard, 2019 Viruses rule over adaptation in conserved human proteins. bioRxiv.

Charlesworth J., and A. Eyre-Walker, 2007 The other side of the nearly neutral theory, evidence of slightly advantageous back-mutations. Proc. Natl. Acad. Sci. U. S. A. 104: 16992–16997.

Cingolani P., A. Platts, L. L. Wang, M. Coon, T. Nguyen, et al., 2012 A program for annotating and predicting the effects of single nucleotide polymorphisms, SnpEff: SNPs in the genome of Drosophila melanogaster strain w1118; iso-2; iso-3. Fly 6: 80–92.

Enard D., L. Cai, C. Gwennap, and D. A. Petrov, 2016 Viruses are a dominant driver of protein adaptation in mammals. Elife 5: e12469.

Eyre-Walker A., M. Woolfit, and T. Phelps, 2006 The distribution of fitness effects of new deleterious amino acid mutations in humans. Genetics 173: 891–900.

Eyre-Walker A., and P. D. Keightley, 2007 The distribution of fitness effects of new mutations. Nat. Rev. Genet. 8: 610–618.

Eyre-Walker A., and P. D. Keightley, 2009 Estimating the rate of adaptive molecular evolution in the presence of slightly deleterious mutations and population size change. Mol. Biol. Evol. 26: 2097–2108.

Fisher R. A., 1930 The Genetical Theory of Natural Selection Oxford Univ. Press (Clarendon), London (Reprinted and revised, 1958).

Galtier N., 2016 Adaptive Protein Evolution in Animals and the Effective Population Size Hypothesis. PLoS Genet. 12: e1005774.

Glémin S., 2003 HOW ARE DELETERIOUS MUTATIONS PURGED? DRIFT VERSUS NONRANDOM MATING. Evolution 57: 2678.

Glémin S., P. F. Arndt, P. W. Messer, D. Petrov, N. Galtier, et al., 2015 Quantification of GC-biased gene conversion in the human genome. Genome Res. 25. https://doi.org/10.1101/gr.185488.114

Gordo I., and F. Dionisio, 2005 Nonequilibrium model for estimating parameters of deleterious mutations. Phys. Rev. E Stat. Nonlin. Soft Matter Phys. 71: 031907.

Gossmann T. I., B.-H. Song, A. J. Windsor, T. Mitchell-Olds, C. J. Dixon, et al., 2010 Genome wide analyses reveal little evidence for adaptive evolution in many plant species. Mol. Biol. Evol. 27: 1822–1832.

Gossmann T. I., P. D. Keightley, and A. Eyre-Walker, 2012 The effect of variation in the effective population size on the rate of adaptive molecular evolution in eukaryotes. Genome Biol. Evol. 4: 658–667.

Grossen C., F. Guillaume, L. F. Keller, and D. Croll, 2019 Accumulation and purging of deleterious mutations through severe bottlenecks in ibex. bioRxiv.

Han S., A. M. Andrés, T. Marques-Bonet, and M. Kuhlwilm, 2019 Genetic variation in Pan species is shaped by demographic history and harbors lineage-specific functions. Genome Biol. Evol. https://doi.org/10.1093/gbe/evz047

Hartl D. L., and C. H. Taubes, 1996 Compensatory nearly neutral mutations: selection without adaptation. J. Theor. Biol. 182: 303–309.

Hernandez R. D., S. H. Williamson, and C. D. Bustamante, 2007 Context dependence, ancestral misidentification, and spurious signatures of natural selection. Mol. Biol. Evol. 24: 1792–1800.

Huber C. D., B. Y. Kim, C. D. Marsden, and K. E. Lohmueller, 2017 Determining the factors driving selective effects of new nonsynonymous mutations. Proc. Natl. Acad. Sci. U. S. A. 114: 4465–4470.

James J., D. Castellano, and A. Eyre-Walker, 2016 DNA sequence diversity and the efficiency of natural selection in animal mitochondrial DNA. Heredity. https://doi.org/10.1038/hdy.2016.108

Jónsson H., P. Sulem, B. Kehr, S. Kristmundsdottir, F. Zink, et al., 2017 Parental influence on human germline de novo mutations in 1,548 trios from Iceland. Nature 549: 519–522.

Keightley P. D., and A. Eyre-Walker, 2007 Joint inference of the distribution of fitness effects of deleterious mutations and population demography based on nucleotide polymorphism frequencies. Genetics 177: 2251–2261.

Keightley P. D., and B. C. Jackson, 2018 Inferring the Probability of the Derived vs. the Ancestral Allelic State at a Polymorphic Site. Genetics 209: 897–906.

Kim B. Y., C. D. Huber, and K. E. Lohmueller, 2017 Inference of the Distribution of Selection Coefficients for New Nonsynonymous Mutations Using Large Samples. Genetics 206: 345–361.

Kimura M., T. Maruyama, and J. F. Crow, 1963 The mutation load in small populations. Genetics. Kimura M., 1983 The Neutral Theory of Molecular Evolution. Cambridge University Press.

Kjeldsen K. U., T. Bataillon, N. Pinel, S. De Mita, M. B. Lund, et al., 2012 Purifying selection and molecular adaptation in the genome of Verminephrobacter, the heritable symbiotic bacteria of earthworms. Genome Biol. Evol. 4: 307–315.

Kousathanas A., and P. D. Keightley, 2013 A comparison of models to infer the distribution of fitness effects of new mutations. Genetics. https://doi.org/10.1534/genetics.112.148023

Kronenberg Z. N., I. T. Fiddes, D. Gordon, S. Murali, S. Cantsilieris, et al., 2018 High-resolution comparative analysis of great ape genomes. Science 360. https://doi.org/10.1126/science.aar6343

Lourenço J. M., S. Glémin, and N. Galtier, 2013 The rate of molecular adaptation in a changing environment. Mol. Biol. Evol. 30: 1292–1301.

Mailund T., J. Y. Dutheil, A. Hobolth, G. Lunter, and M. H. Schierup, 2011 Estimating divergence time and ancestral effective population size of Bornean and Sumatran orangutan subspecies using a coalescent hidden Markov model. PLoS Genet. 7: e1001319.

Manuel M. de, M. Kuhlwilm, P. Frandsen, V. C. Sousa, T. Desai, et al., 2016 Chimpanzee genomic diversity reveals ancient admixture with bonobos. Science 354: 477–481.

McManus K. F., J. L. Kelley, S. Song, K. R. Veeramah, A. E. Woerner, et al., 2015 Inference of gorilla demographic and selective history from whole-genome sequence data. Mol. Biol. Evol. 32: 600–612.

Messer P. W., and D. a. Petrov, 2013 Frequent adaptation and the McDonald-Kreitman test. Proc. Natl. Acad. Sci. U. S. A. 110: 8615–8620.

Nam K., K. Munch, T. Mailund, A. Nater, M. P. Greminger, et al., 2017 Evidence that the rate of strong selective sweeps increases with population size in the great apes. Proc. Natl. Acad. Sci. U. S. A. 114: 1613–1618.

Ohta T., 1992 The Nearly Neutral Theory of Molecular Evolution. Annu. Rev. Ecol. Syst. 23: 263–286.

Pagel M., and A. Meade, 2006 Bayesian analysis of correlated evolution of discrete characters by reversible-jump Markov chain Monte Carlo. Am. Nat. 167: 808–825.

Phifer-Rixey M., F. Bonhomme, P. Boursot, G. A. Churchill, J. Piálek, et al., 2012 Adaptive evolution and effective population size in wild house mice. Mol. Biol. Evol. 29: 2949–2955.

Poon A., and S. P. Otto, 2000 Compensating for our load of mutations: freezing the meltdown of small populations. Evolution 54: 1467–1479.

Pouyet F., S. Aeschbacher, A. Thiéry, and L. Excoffier, 2018 Background selection and biased gene conversion affect more than 95% of the human genome and bias demographic inferences. Elife 7. https://doi.org/10.7554/eLife.36317

Prado-Martinez J., P. H. Sudmant, J. M. Kidd, H. Li, J. L. Kelley, et al., 2013 Great ape genetic diversity and population history. Nature 499: 471–475.

Rousselle M., M. Mollion, B. Nabholz, T. Bataillon, and N. Galtier, 2018 Overestimation of the adaptive substitution rate in fluctuating populations. Biol. Lett. 14. https://doi.org/10.1098/rsbl.2018.0055

Rousselle M., P. Simion, M. K. Tilak, E. Figuet, B. Nabholz, et al., 2019 Is adaptation limited by mutation? A timescale dependent effect of genetic diversity on the adaptive substitution rate in animals. bioRxiv 643619.

Scally A., J. Y. Dutheil, L. W. Hillier, G. E. Jordan, I. Goodhead, et al., 2012 Insights into hominid evolution from the gorilla genome sequence. Nature 483: 169–175.

Schneider A., B. Charlesworth, A. Eyre-Walker, and P. D. Keightley, 2011 A method for inferring the rate of occurrence and fitness effects of advantageous mutations. Genetics 189: 1427–1437.

Ségurel L., M. J. Wyman, and M. Przeworski, 2014 Determinants of mutation rate variation in the human germline. Annu. Rev. Genomics Hum. Genet. 15: 47–70.

Silander O. K., O. Tenaillon, and L. Chao, 2007 Understanding the evolutionary fate of finite populations: the dynamics of mutational effects. PLoS Biol. 5: e94.

Smith T. C. A., P. F. Arndt, and A. Eyre-Walker, 2018 Large scale variation in the rate of germ-line de novo mutation, base composition, divergence and diversity in humans. PLoS Genet. 14: e1007254.

Stevison L. S., A. E. Woerner, J. M. Kidd, J. L. Kelley, K. R. Veeramah, et al., 2016 The Time Scale of Recombination Rate Evolution in Great Apes. Mol. Biol. Evol. 33: 928–945.

Strasburg J. L., N. C. Kane, A. R. Raduski, A. Bonin, R. Michelmore, et al., 2011 Effective population size is positively correlated with levels of adaptive divergence among annual sunflowers. Mol. Biol. Evol. 28: 1569–1580.

Tataru P., M. Mollion, S. Glémin, and T. Bataillon, 2017 Inference of Distribution of Fitness Effects and Proportion of Adaptive Substitutions from Polymorphism Data. Genetics 207: 1103–1119.

Tataru P., and T. Bataillon, 2019 polyDFEv2.0: Testing for invariance of the distribution of fitness effects within and across species. Bioinformatics. https://doi.org/10.1093/bioinformatics/bty1060

Thomas G. W. C., R. J. Wang, A. Puri, R. A. Harris, M. Raveendran, et al., 2018 Reproductive Longevity Predicts Mutation Rates in Primates. Curr. Biol. 28: 3193–3197.e5.

Torres R., M. G. Stetter, R. D. Hernandez, and J. Ross-Ibarra, 2019 The temporal dynamics of background selection in non-equilibrium populations. BioRxiv.

Uricchio L. H., D. A. Petrov, and D. Enard, 2019 Exploiting selection at linked sites to infer the rate and strength of adaptation. Nat Ecol Evol. https://doi.org/10.1038/s41559-019-0890-6

Williamson S. H., R. Hernandez, A. Fledel-Alon, L. Zhu, R. Nielsen, et al., 2005 Simultaneous inference of selection and population growth from patterns of variation in the human genome. Proc. Natl. Acad. Sci. U. S. A. 102: 7882–7887.

Zhen Y., C. D. Huber, R. W. Davies, and K. E. Lohmueller, 2018 Stronger and higher proportion of beneficial amino acid changing mutations in humans compared to mice and flies. bioRxiv 427583.

